# Partial FAM19A5 Deficiency in Mice Leads to Disrupted Spine Maturation, Hyperactivity, and an Altered Fear Response

**DOI:** 10.1101/2024.04.29.591582

**Authors:** Anu Shahapal, Sumi Park, Sangjin Yoo, Jong-Ik Hwang, Jae Young Seong

## Abstract

The FAM19A*5* polypeptide, encoded by the TAFA*5* gene, is evolutionarily conserved among vertebral species. This protein is predominantly expressed in the brain, highlighting its crucial role in the central nervous system. Here, we investigated the potential roles of FAM19A*5* in brain development and behavior using a FAM19A*5*-LacZ KI mouse model. This model exhibited a partial reduction in the FAM19A*5* protein level. FAM19A*5*-LacZ KI mice displayed no significant alterations in gross brain structure but alterations in dendritic spine distribution, with a bias toward immature forms. These mice also had lower body weights. Behavioral tests revealed that compared with their wild-type littermates, FAM19A*5*-LacZ KI mice displayed hyperactivity and a delayed innate fear response. These findings suggest that FAM19A*5* plays a role in regulating spine formation and maintenance, thereby contributing to neural connectivity and behavior.

## INTRODUCTION

FAM19A*5*, a polypeptide encoded by the TAFA*5* gene located on chromosome 22 (22q13.32) in humans and chromosome 15 in mice, is primarily expressed in the brain^1^. This restricted expression and the highly conserved amino acid sequence across vertebrates suggest a crucial role for FAM19A*5* in brain development and function^2,3^. A study by Huang et al. showed that deletion of FAM19A*5* led to reduced spine density, glutamate signaling and neuronal activity. These changes resulted in depressive-like behavior and impaired spatial memory in mice, suggesting that FAM19A*5* acts as a key regulator of early brain development and cognitive function^4^. Further strengthening this connection, a recent study demonstrated that targeting FAM19A*5* with an antibody reversed synaptic loss and improved cognitive function in mouse models of Alzheimer’s disease, suggesting its potential as a therapeutic target^5^. In addition, there is a report showing a potential role for FAM19A1, a FAM19A*5* paralog, in regulating food intake and metabolism. Changes in FAM19A1 mRNA levels in brain regions such as the hypothalamus, cerebral cortex, and cerebellum were detected when the feeding state of the mice was altered^6^.

Additionally, a few human cases of developmental defects associated with deletions in chromosome 22, where FAM19A*5* is located, have been reported^7,8^. The microdeletion of chromosome 22, del22q13.32-q13.33, encompasses exon 4 of FAM19A*5* and is accompanied by clinical features such as hyperactivity, aggression, low body weight, skeletal abnormalities, speech disorders, and brain deformities. This finding suggested that loss of FAM19A*5* function disrupts critical aspects of brain development and function, potentially contributing to a range of behavioral and cognitive impairments.

Here, we investigated the potential role of FAM19A*5* in the brain using a FAM19A*5*-LacZ KI mouse model^3,9^. LacZ includes its own poly A-tail and stop codon, which were integrated in the upstream of exon 4 of FAM19A*5*, preventing the formation of complete FAM19A*5* mRNA and subsequent production of functional FAM19A*5* protein. Western blotting confirmed reduced FAM19A*5* expression in FAM19A*5*-LacZ KI mice, validating this model as a partial knockout (KO) model of FAM19A*5*. Any physical deformities and behavioral changes in FAM19A*5*-LacZ KI mice after a reduction in the FAM19A*5* level were investigated to determine the role of FAM19A*5* in the regulation of brain functions.

## RESULTS

### FAM19A5-LacZ KI mice as a partial FAM19A5 KO mouse model

To generate a partial knockout FAM19A*5* mouse model, we integrated LacZ upstream of exon 4 of the FAM19A*5*. Western blot analysis of brain lysates from FAM19A*5*-LacZ KI mice (FAM19A*5*^+/LacZ^ and FAM19A*5*^LacZ/LacZ^ genotypes) revealed significant reductions in FAM19A*5* protein levels compared to those in wild type mice (WT, FAM19A*5*^+/+^) (Figure 1A and 1B). This finding confirms that the FAM19A*5*-LacZ KI model effectively reduces FAM19A*5* protein expression and is suitable for studying the function of FAM19A*5* in vivo.

**Fig. 1.**
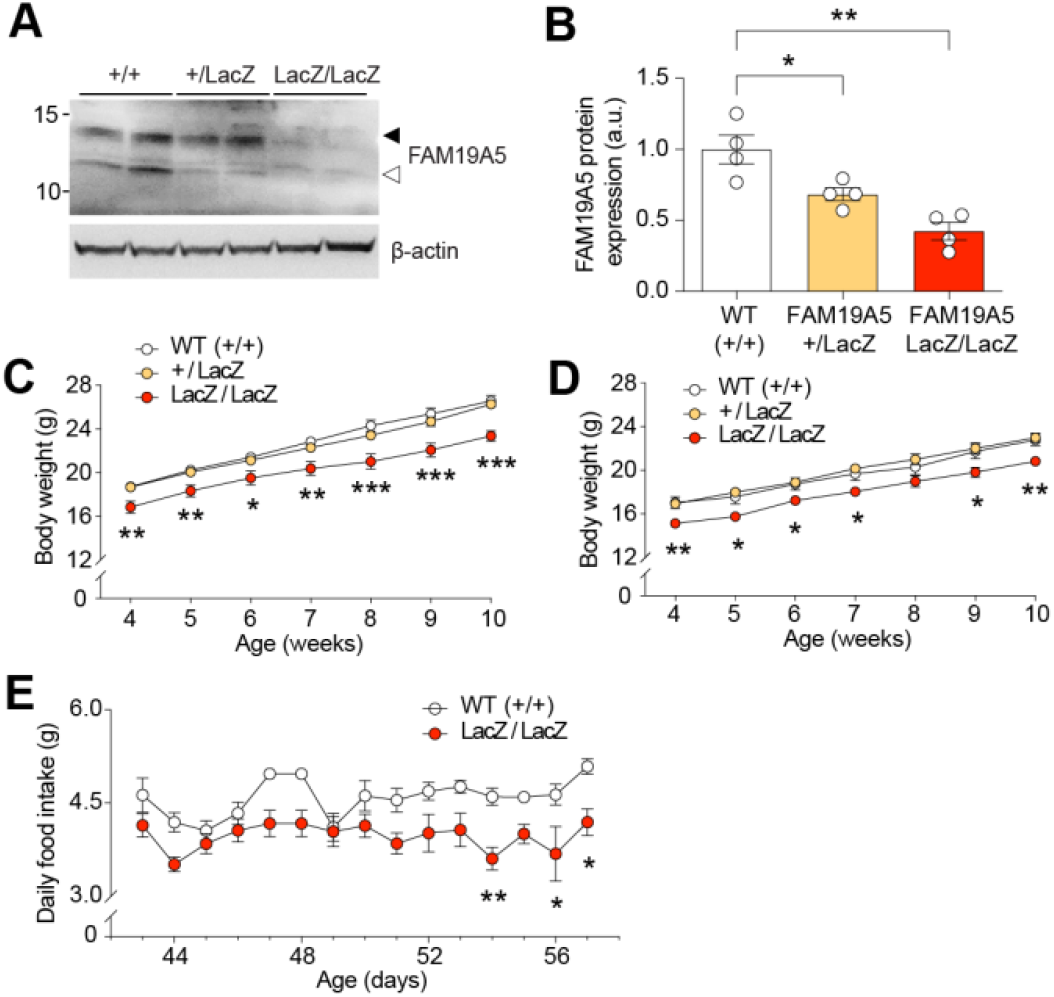
Reduction of FAM19A*5* protein levels in FAM19A*5*-LacZ KI mice. (A) Western blot analysis of FAM19A*5* protein in FAM19A*5*-LacZ KI mice and WT littermates, solid arrow: glycosylated FAM19A*5*, open arrow: nonglycosylated FAM19A*5*. (B) FAM19A*5* protein expression level in the brain of FAM19A*5*^+/LacZ^, FAM19A*5*^LacZ/LacZ^, and FAM19A*5*^+/+^ littermates, n=3, One way ANOVA followed by bonferroni’s multiple comparisons test. Reduced body weight in FAM19A*5*^LacZ/LacZ^, both (C) male and (D) female mice compared to FAM19A*5*^+/+^ littermates, male: n=16 (WT^+/+^), 27 (FAM19A*5*^+/LacZ^), 15 (FAM19A*5*^LacZ/LacZ^), female: n=11 (WT^+/+^), 18 (FAM19A*5*^+/LacZ^), 11 (FAM19A*5*^LacZ/LacZ^). Two way ANOVA followed by bonferroni’s multiple comparisons test (E) Reduced daily food intake in male FAM19A*5*^LacZ/LacZ^ and WT^+/+^ mice for 15 days, n=7 (WT^+/+^), 6 (FAM19A*5*^LacZ/LacZ^). Data are presented as the mean ± SEM. Two way ANOVA followed by bonferroni’s multiple comparisons test, *P<0.05; **P<0.01 and ***P<0.001 vs. WT^+/+^.

### Reduced body weight and food intake in FAM19A5-LacZ KI mice

Since FAM19A*5* expression begins as early as embryonic day 10.5^3^, a reduction in FAM19A*5* expression throughout development in FAM19A*5*-LacZ KI mice might cause abnormalities. When heterozygous male and female mice were mated to generate homozygous mice, 19.92% (22.737% male and 17.054% female) of the FAM19A*5*^LacZ/LacZ^ mice were born, which was lower than the expected 25%. Importantly, these mice displayed similar sex ratios with no visible physical deformities and displayed normal postnatal growth and survival compared to their FAM19A*5*^+/+^ littermates. Despite normal growth and survival, FAM19A*5*-LacZ mice exhibited significantly lower weight compared to FAM19A*5*^+/+^ mice (Figure 1C, D). In addition, daily food intake was relatively lower in the FAM19A*5*^LacZ/LacZ^ mice than in the FAM19A*5*^+/+^ mice, with significant decreases on the 54th, 56th, and 57th days of age (Figure 1E).

### Gross brain morphology in FAM19A5-LacZ KI mice

Given its early expression and predominant expression in neuronal stem cells (NSCs) and progenitor cells within the brain of germinal zones^3^, a partial reduction in FAM19A*5* may affect normal brain development and growth. To investigate this possibility, whole-brain size was measured in adult FAM19A*5*-LacZ KI mice and compared with WT mice. There were no significant differences in total brain length, width, and the cortical length between the FAM19A*5*-LacZ KI mice and their WT littermates (Figure 2A-C).

**Fig. 2.**
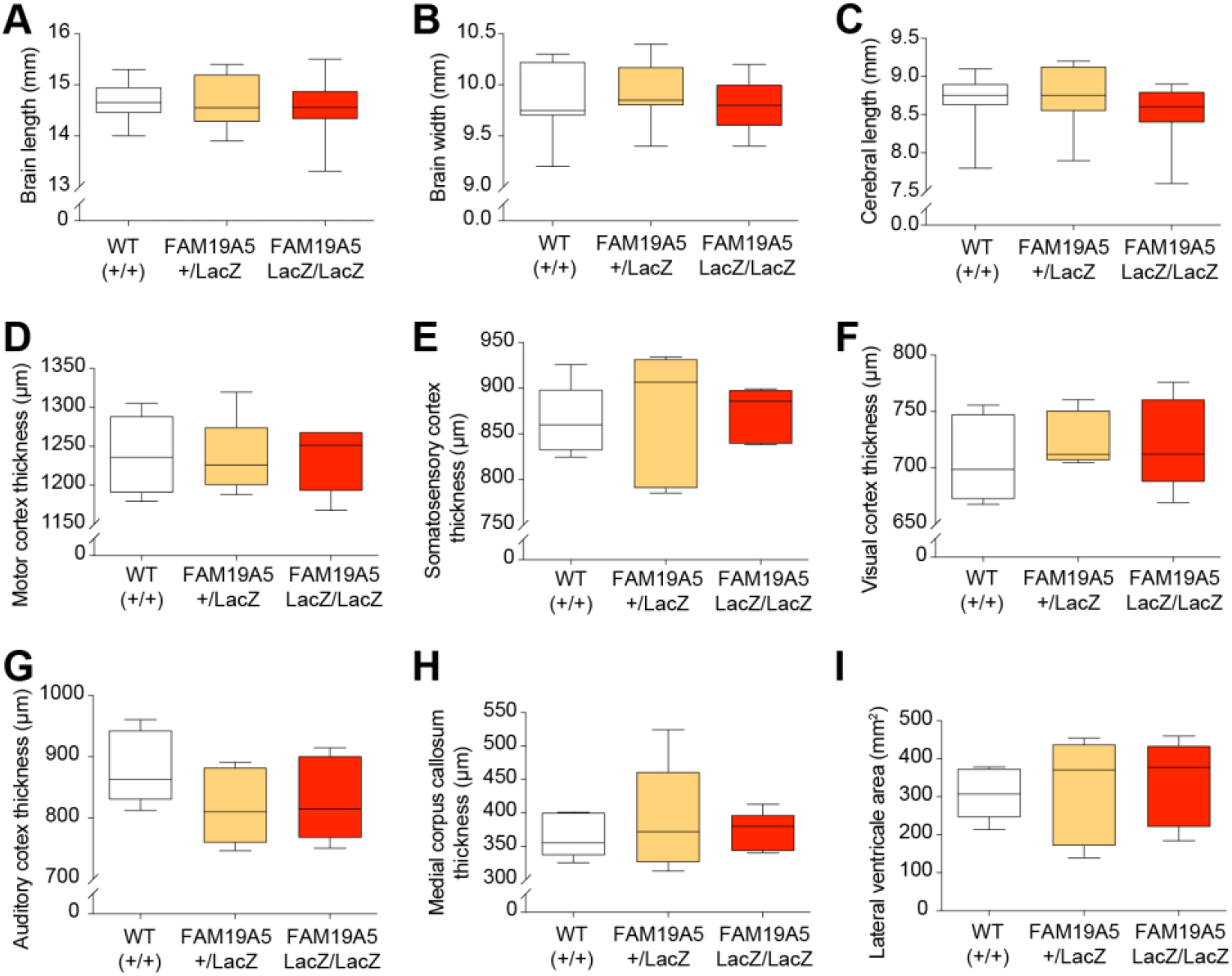
A comparison of brain structure between FAM19A*5*-LacZ KI mice and wild-type littermates. (A) Whole brain length, (B) width, and (C) cerebral cortical length measurement between FAM19A*5*-LacZ KI mice and FAM19A*5*^+/+^ littermates. n=14 (WT^+/+^), 14 (FAM19A*5*^+/LacZ^) and 16 (FAM19A*5*^LacZ/LacZ^). Thickness of (D) motor, (E) somatosensory, (F) visual, and (G) auditory cortex between the mice (n=5). Thickness of (H) medial corpus callosum and (I) area of lateral ventricle between the mice (n=5 each).

To assess potential effects on cortical development, we performed detailed histological measurements. The thicknesses of various cortical regions, including the motor, somatosensory, visual, and auditory cortex were measured in Nissl-stained coronal brain sections. FAM19A*5*-LacZ KI mice exhibited normal cortical thickness (Figure 2D-G). Notably, some FAM19A*5*-LacZ KI mice displayed enlarged lateral ventricles (LVs) compared to those of WT mice. Hence, the area of the LV and thickness of the corpus callosum were measured, but no significant differences were found compared to those of the WT group (Figure 2H and I). Furthermore, the thicknesses of cortical layers 1, 2/3, 4, *5*, and 6 were compared between FAM19A*5*-LacZ KI and WT mice. No significant differences were observed. (Supplementary Figure 1). These histological analyses revealed normal gross brain morphology in FAM19A*5*-LacZ KI mice, suggesting that a partial reduction in FAM19A*5* may not significantly impact overall brain development.

### Dendritic spine analysis in FAM19A5-LacZ KI mice

Building on previous findings that a complete FAM19A*5* knockout reduces spine density^4^, we hypothesized that a partial FAM19A*5* knockout mouse model may also exhibit a decrease in spine density. Spine density and length-to-width (LWR) were analyzed in both basal and apical dendritic spines of pyramidal neurons from layers 2/3 and *5* of the motor and somatosensory cortex using Golgi staining.

In the motor cortex, the spine density was not significantly altered, both in apical and basal dendrites of layers 2/3 and *5* between the FAM19A*5*^LacZ/LacZ^ mice and their FAM19A*5*^+/+^ littermates (Figure 3A, B, D, E). However, compared with their FAM19A*5*^+/+^ littermates, FAM19A*5*^LacZ/LacZ^ mice exhibited a significant increase in the LWR of both apical and basal dendrites in layer *5* pyramidal neurons (Figure 3F). Although the LWR of the apical and basal dendritic spines of layer 2/3 neurons were not significantly different, a trend toward an increase was observed in the FAM19A*5*^LacZ/LacZ^ group compared to that in the FAM19A*5*^+/+^ littermates (Figure 3C).

**Fig. 3.**
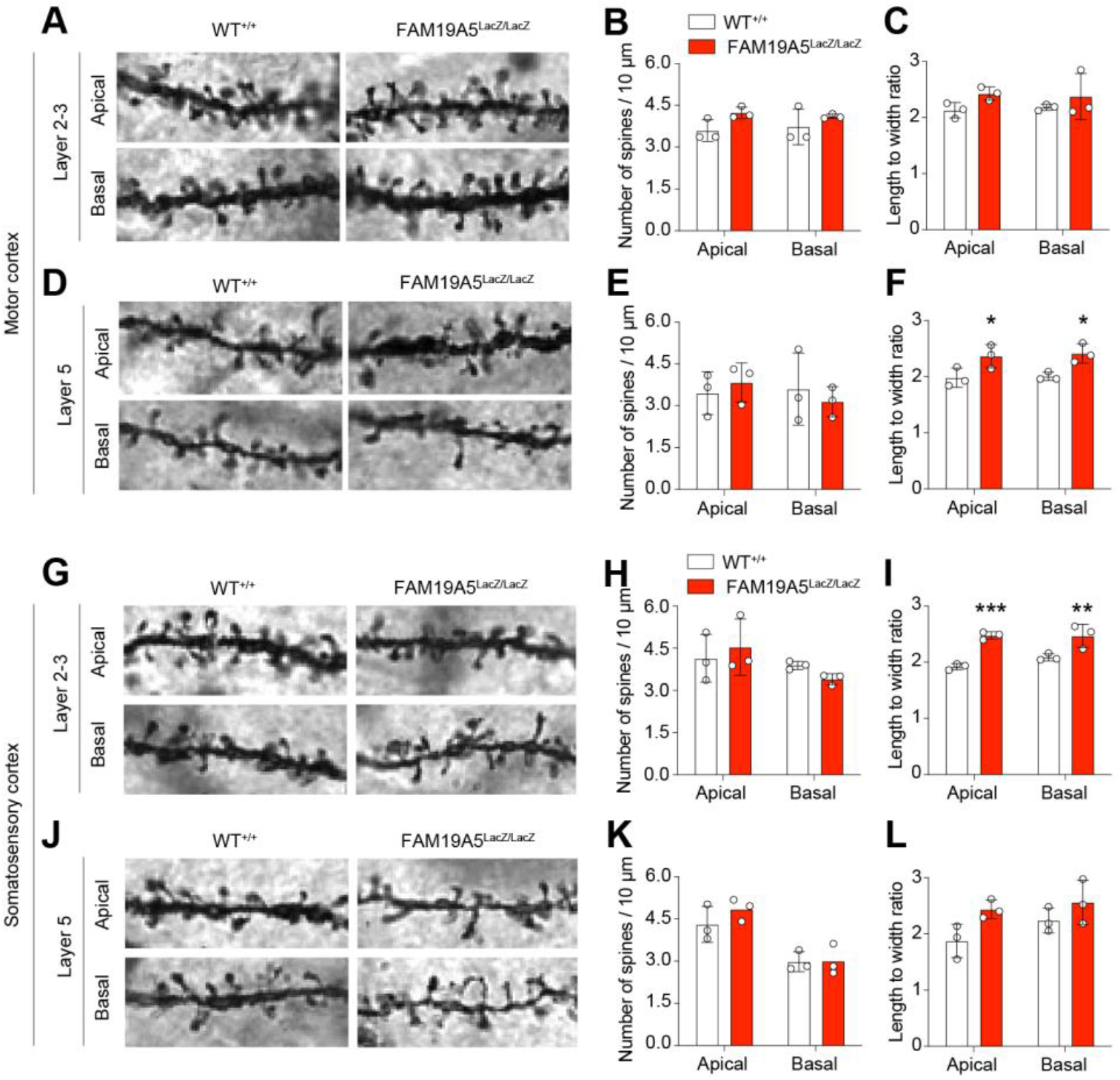
Dendritic spine analysis in the adult FAM19A*5*-LacZ KI mouse brain. Spine density and length-width ratio of apical and basal dendritic spines in layer 2/3 and *5* pyramidal neurons of the motor cortex and somatosensory cortex. (A) Golgi staining image of dendritic spine in layer 2/3 of the motor cortex. And its quantification in terms of (B) density and (C) length-width ratio. (D) Golgi staining image of dendritic spine in layer *5* of the motor cortex. And its quantification in terms of (E) density and (F) length-width ratio. (G) Golgi staining image of dendritic spine in layer 2/3 of the somatosensory cortex. And its quantification in terms of (H) density and (I) length-width ratio. (J) Golgi staining image of dendritic spine in layer *5* of the motor cortex. And its quantification in terms of (K) density and (L) length-width ratio. Data are presented as the mean ± SEM. n=3 each, Two way ANOVA followed by bonferroni’s multiple comparisons test, *P<0.05, **P<0.01 and ***P<0.001 vs. WT^+/+^.

Similarly, in the somatosensory cortex, spine density remained unchanged between FAM19A*5*^LacZ/LacZ^ and FAM19A*5*^+/+^ mice (Figure 3G, H, J, K). However, FAM19A*5*^LacZ/LacZ^ mice displayed a significant increase in the LWR of both apical and basal dendritic spines in layers 2/3 compared to FAM19A*5*^+/+^ littermates (Figure 3I). Although the LWR of basal dendritic spines in layer *5* was not significantly altered, a trend towards an increase was observed in FAM19A*5*^LacZ/LacZ^ mice (Figure 3L).

Morphologically, dendritic spines are categorized into various types. Filopodia and thin, characterized by their long, motile structure, are considered immature and unstable spine forms. In contrast, mature and stable spines, known as mushroom spines, are typically shorter and wider in size^10^. The observed increase in the LWR of spines in FAM19A*5*^LacZ/LacZ^ mice suggested a potential increase in the number of immature dendritic spines in these adult mice.

To test this hypothesis, we classified the spines into four types; filopodia, thin, branched, and mushroom spine. In the motor cortex, FAM19A*5*^LacZ/LacZ^ mice displayed a trend toward increased filopodia, thin and branched spines, along with a decrease in mushroom spines, in both the apical and basal dendrites of layers 2/3 and 5, although these changes did not reach statistical significance compared to those of the FAM19A*5*^+/+^ controls (Figure 4A-D). The somatosensory cortex mirrored this trend, with FAM19A*5*^LacZ/LacZ^ mice exhibiting a significant increase in filopodia and thin, and a decrease in mushroom spines in both the apical and basal dendrites of layer 2/3 neurons compared to those of FAM19A*5*^+/+^ controls (Figure 4E, F). In addition, the number of branched spines, which are also considered functionally unstable spines, tended to increase in FAM19A*5*^LacZ/LacZ^ mice. In layer *5* of the somatosensory cortex, although the differences were not statistically significant, both apical and basal dendrites tended to exhibit increased filopodia, thin and branched spines and decreased mushroom spines in the brains of FAM19A*5*^LacZ/LacZ^ mice compared to those in the brains of FAM19A*5*^+/+^ mice (Figure 4G, H). The morphology of spines is closely associated with their function and synaptic strength^11,12^. The observed shift toward immature spine types in FAM19A*5*-deficient mice suggests a potential disruption in synaptic plasticity, which may underlie the reduced body weight and daily food intake observed in these animals.

**Fig. 4.**
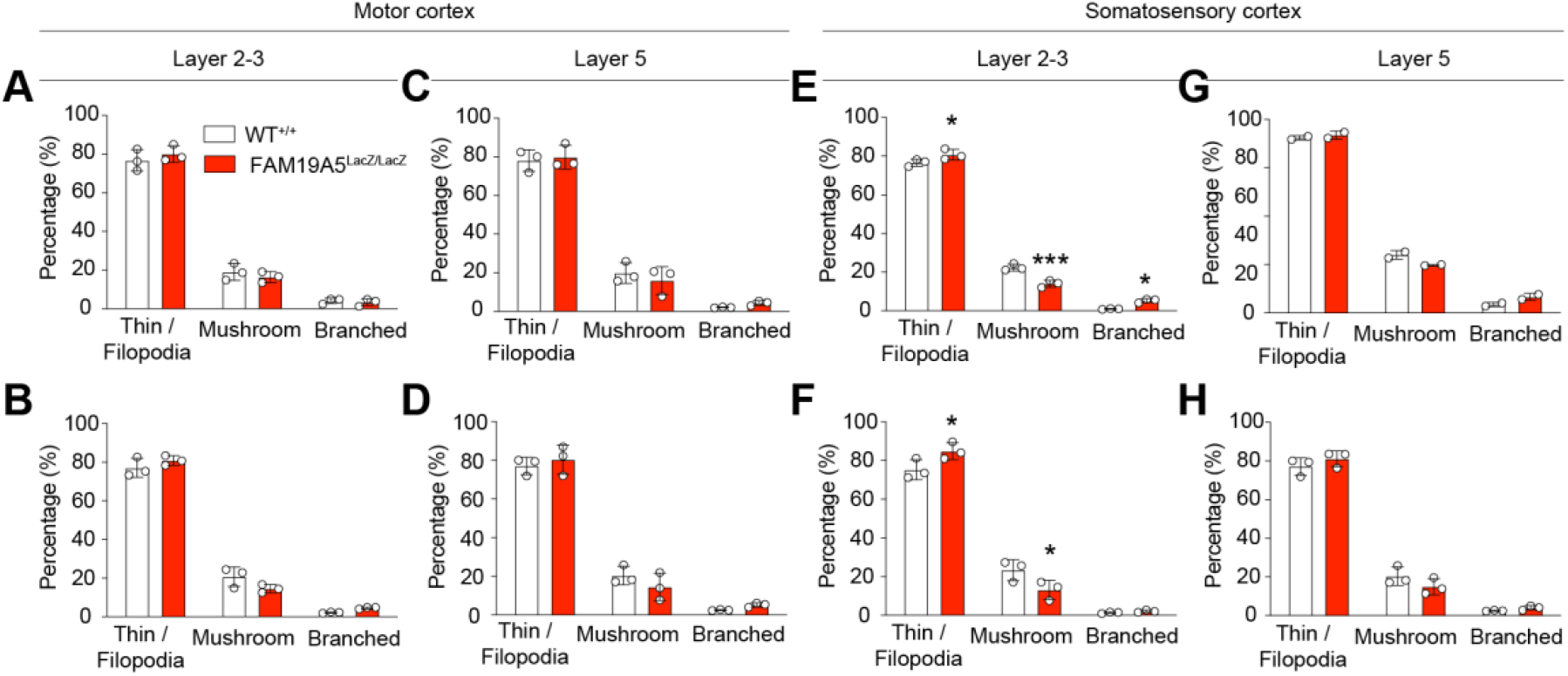
Alteration in dendritic spine type in the adult FAM19A*5*-LacZ KI mouse brain. Classified spine types in (A) apical and (B) basal dendrites of layer 2/3 pyramidal neurons in motor cortex. Classification of spine types in (C) Apical and (D) basal dendrites of layer *5* pyramidal neurons in motor cortex. Classified spine types in (E) apical and (F) basal dendrites of layer 2/3 pyramidal neurons in motor cortex. Classification of spine types in (G) Apical and (H) basal dendrites of layer *5* pyramidal neurons in motor cortex. Data are presented as the mean ± SEM. n=3 (WT), 2 (FAM19A*5*^+/LacZ^) and n=3 (FAM19A*5*^LacZ/LacZ^), *P<0.05, ***P<0.001 vs. WT.

### General locomotion, exploratory and motor function in FAM19A5-LacZ KI mice

The observed shift in immature spines and decrease in mature spines in FAM19A*5*-LacZ KI mice, which are indicative of altered synaptic function and plasticity, may disrupt normal brain function and contribute to behavioral abnormalities. To investigate this possibility, we performed a series of behavioral tests. General locomotor, exploratory, and anxiety-related activities were observed using an open field test (OFT). The results showed a significant increase in locomotor activity in the FAM19A*5*^LacZ/LacZ^ mice compared to that in the FAM19A*5*^+/+^mice, as indicated by the total distance traveled and mean speed of movement (Figure *5*A-C). This hyperactivity was primarily observed in the peripheral region of the open field but not in the central zone (Figure *5*D-H). Notably, there were no significant differences in the time spent in the center zone (Figure *5*I), suggesting an absence of anxiety-like behavior in FAM19A*5*-LacZ KI mice compared to WT littermates.

**Fig. 5.**
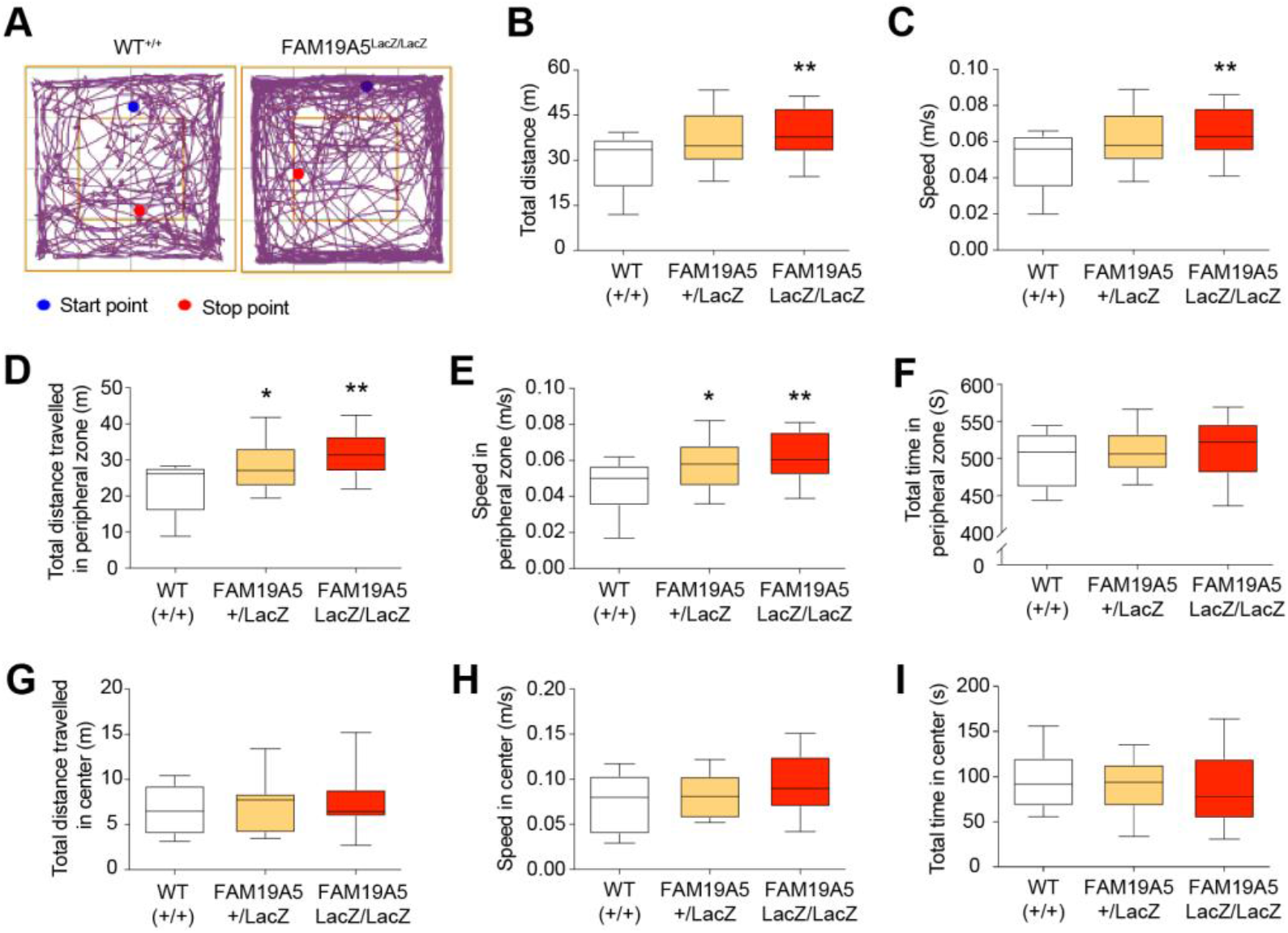
Hyperactivity in FAM19A*5*-LacZ KI mice. (A) Representative track plot of movements in open field box during 10 min of exploration time in FAM19A*5*^LacZ/LacZ^ mice and FAM19A*5*^+/+^ littermates. (B) Total traveling distance and (C) mean speed of movement in the open field, respectively. (D) Total traveling distance, (E) mean speed of movement and (F) time spent in the peripheral zone of the open field box, respectively. (G) Total traveling distance, (H) mean speed movement and (I) time spent in the center region of the open field, respectively. n=13 (WT^+/+^), n=11 (FAM19A*5*^+/LacZ^), n=16 (FAM19A*5*^LacZ/LacZ^). Data are presented as the mean ± SEM. *P<0.05 **P<0.01 vs. FAM19A*5*^+/+^.

Rearing, where mice stand upright on their hind limbs, is considered a measure of exploratory behavior. There are two types of rearing: supported rearing, where mice use the arena wall for balance, which is indicative of enhanced locomotor activity, and unsupported rearing, which reflects an emotional state^13,14^. Although there was no significant difference in total rearing number (Supplementary Figure 2A), FAM19A*5*^LacZ/LacZ^ mice displayed a significant increase in supported rearing accompanied by a decrease in unsupported rearing compared to that of the FAM19A*5*^+/+^ group (Supplementary Figure 2B and C). These findings suggest a potential dampening of the emotional component of exploration in FAM19A*5*-LacZ KI mice. This shift, with increased supported rearing and decreased unsupported rearing, might indicate a reduced anxiety-like behavior in these mice compared to their wild-type littermates.

In addition to rearing, grooming was analyzed to assess repetitive, compulsive-like behavior. Although there was no significant difference in total grooming time, compared with FAM19A*5*^+/+^ mice, FAM19A*5*^LacZ/LacZ^ mice tended to exhibit decreased grooming (Supplementary Figure 2D), suggesting the absence of repetitive-like behavior.

Next, we performed the rotarod test and hanging wire test to evaluate motor learning, coordination, and strength in FAM19A*5*-LacZ KI mice. FAM19A*5*^LacZ/LacZ^ mice initially tended to fall sooner than did FAM19A*5*^+/+^ control mice in the first two days (Supplementary Figure 2E), but their performance recovered to similar levels by the later sessions. In the hanging wire test, compared with the FAM19A*5*^+/+^ mice, the FAM19A*5*^LacZ/LacZ^ mice tended to fall (Supplementary Figure 2F), but the difference in the number of falls was not statistically significant. Overall, these results suggest normal motor function in FAM19A*5*-LacZ KI mice.

### Repetitive and compulsive-like behavior in FAM19A5-LacZ KI mice

Most neurodevelopmental disorders characterized by hyperactivity, such as attention-deficit/hyperactivity disorder (ADHD) and autism spectrum disorder (ASD), are accompanied by repetitive or compulsive-like behavior. In the OFT, FAM19A*5*-LacZ KI mice tended to exhibit reduced grooming time, suggesting an absence of repetitive-like behavior. To further investigate repetitive behavior in hyperactive FAM19A*5*-LacZ KI mice, the marble burying test and nestlet shredding test were performed^15^. FAM19A*5*^LacZ/LacZ^ mice showed a significant decrease in the number of buried marbles compared to that of FAM19A*5*^+/+^ mice (Supplementary Figure 2G), suggesting reduced repetitive behavior in the FAM19A*5*-deficient mice. Although no significant difference was detected in the nestlet shredding test (Figure Supplementary 2H), these findings collectively indicate the absence of repetitive or compulsive-like behavior in FAM19A*5*-LacZ KI mice.

### Anxiety/depressive-like behavior in FAM19A5-LacZ KI mice

To further confirm the anxiety-related behavior of FAM19A*5*-LacZ KI mice, the elevated plus maze test (EPMT) was performed. Compared with control mice, FAM19A*5*^LacZ/LacZ^ mice showed a significant increase in distance traveled and speed of movement in the maze (Supplementary Figure 3A-C). Although the total time spent and total number of entries in the open arm of the maze were greater in FAM19A*5*^LacZ/LacZ^ mice than in FAM19A*5*^+/+^mice, the differences were not statistically significant (Supplementary Figure 3D and E), suggesting the absence of anxiety-like behavior in FAM19A*5*-LacZ KI mice.

The tail suspension test (TST) was performed to assess depressive-like behavior in FAM19A*5*-LacZ KI mice. As shown in Supplementary Figure 3F, immobility did not significantly differ between the FAM19A*5*-LacZ KI and WT groups, indicating the absence of depressive-like behavior in FAM19A*5*-LacZ KI mice.

### Learning and memory function in FAM19A5-LacZ KI mice

To investigate whether the increase in immature spines in FAM19A*5*-LacZ KI mice leads to impaired learning and memory function, we performed the Y-maze test for spatial working memory, and the novel object recognition (NOR) test for recognition memory, respectively. In the Y-maze test, no significant difference in spontaneous alterations was observed between FAM19A*5*-LacZ KI and WT mice (Supplementary Figure 4A). This finding suggested normal spatial working memory in the FAM19A*5*-deficient mice. However, consistent with the OFT and EPMT results, FAM19A*5*-LacZ KI mice traveled a significantly greater distance and moved at a faster speed within the maze (Supplementary Figure 4B and C). The observed increase in locomotion and exploration within the Y-maze by FAM19A*5*-deficient mice might be attributable to their reduced anxiety-like behavior observed in previous OFT and EPMT test.

In the NOR test, there were no significant differences in preference for the novel object after 6 h (short-term memory) (Supplementary Figure 4D) or after 24 h (long-term memory) (Supplementary Figure 4E) between FAM19A*5*-LacZ KI and WT mice. These results indicated normal recognition memory, both short-term and long-term, in FAM19A*5*-LacZ KI mice.

### Fear response memory and innate fear response in FAM19A5-LacZ KI mice

We then investigated whether FAM19A*5*-deficient mice have deficits in fear response memory consolidation. To investigate this, a Pavlovian fear conditioning test was performed. FAM19A*5*^LacZ/LacZ^ mice exhibited significantly less freezing behavior than did their FAM19A*5*^+/+^ littermates during the initial fear acquisition period (Figure 6A). While all mice displayed increased freezing on the conditioning day, compared with their FAM19A*5*^+/+^ littermates, the FAM19A*5*^LacZ/LacZ^ mice failed to show robust freezing behavior indicative of fear consolidation. Consequently, FAM19A*5*-LacZ KI mice exhibited a significantly reduced freezing rate during both the contextual (Figure 6B) and auditory fear memory tests (Figure 6C). These findings suggest potential deficits in fear response memory consolidation in FAM19A*5*-LacZ KI mice.

**Fig. 6.**
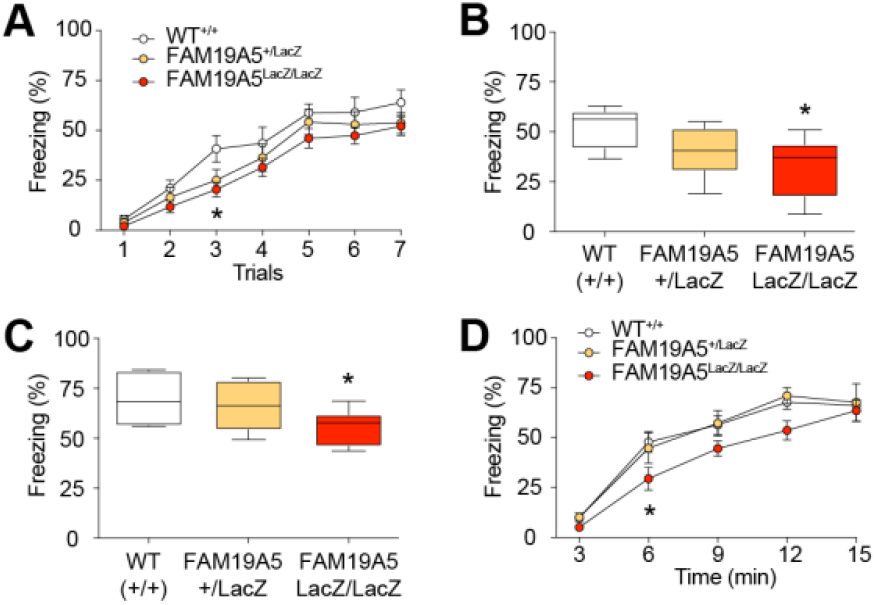
Fear response in FAM19A*5*-LacZ KI mice. (A) Freezing rates were measured in individual trials during the acquisition session in FAM19A*5*-LacZ KI mice and their littermates. The session consisted of 7 repetitive tones (5KHz, 70 dB, 30 s) followed by foot shock (0.7mA, 2 s). Freezing rate during (B) contextual fear memory test and (C) auditory fear memory test. n=7 (WT^+/+^), n=7 (FAM19A*5*^+/LacZ^) and n=7 (FAM19A*5*^LacZ/LacZ^). (D) Freezing rate in response to 2,5-dihydro-2,4,5-trimethylthiazoline exposure was measured in FAM19A*5*-LacZ KI mice and their littermates to test innate fear response. n=15 (WT^+/+^), 4 (FAM19A*5*^+/LacZ^) and 11 (FAM19A*5*^LacZ/LacZ^). Data are presented as the mean ± SEM. *P<0.05 vs. WT^+/+^.

The reduction in freezing rate during the fear acquisition phase in the FAM19A*5*^LacZ/LacZ^ group might be due to their hyperactivity. To investigate whether impaired fear acquisition is related to a general deficit in the fear response, the innate fear response was tested using threat match tracking (TMT). Normally, when mice encounter predators, they tend to exhibit instinctive behaviors such as freezing as a measure of fear.

Upon exposure to the TMT, FAM19A*5*^LacZ/LacZ^ mice exhibited a significant decrease in freezing time compared to FAM19A*5*^+/+^ mice. These mice displayed a significant decrease in freezing time throughout the initial 6 mins of TMT exposure. However, their freezing behavior gradually increased over time, eventually reaching levels similar to those of their FAM19A*5*^+/+^ littermates (Figure 6D). This, coupled with their hyperactivity, may contribute to the impaired fear memory consolidation observed in these animals.

## DISCUSSION

In this study, FAM19A*5*-LacZ KI mice were used as a partial FAM19A*5* KO mouse model to investigate loss-of-function studies of FAM19A*5*. Western blot analysis confirmed a partial reduction in the FAM19A*5* protein in the adult brains of FAM19A*5*-LacZ KI mice compared to WT mice. General physical observation revealed no visible differences in gross appearance and survival among the FAM19A*5*-LacZ KI mice. However, both male and female FAM19A*5*^LacZ/LacZ^ mice had low body weights, which is consistent with the findings of a recent study using a complete FAM19A*5* knockout mouse model^**4**^. Furthermore, compared with the FAM19A*5*^+/+^ mice, the FAM19A*5*^LacZ/LacZ^ mice consumed relatively less food. Additionally, a study reported that alterations in FAM19A*5* expression in the brain depend on nutritional status^**6**^. They observed increased FAM19A*5* mRNA levels in the hypothalamus, cortex, and hippocampus of fasted mice. Notably, the hypothalamus is a brain region known to regulate feeding and satiety and body metabolism^**16**^. Moreover, X-gal staining of adult mouse brains revealed widespread FAM19A*5* expression in the hypothalamus, which was further confirmed by single-cell RNA sequencing results from www.mousebrain.org showing high FAM19A*5* expression in the NPY/AgRP neurons of the arcuate hypothalamic nucleus. These neurons are known as orexigenic neurons, the activation of which results in the induction of food intake^**17**^. Therefore, the partial reduction in FAM19A*5* in FAM19A*5*-LacZ KI mice likely contributes to their lower body weight through a combination of decreased food intake and potentially increased energy expenditure due to their hyperactive behavior.

FAM19A*5*-LacZ KI mice exhibited a normal brain size, as assessed by gross observation. Histological analysis of adult FAM19A*5*-LacZ KI brain sections revealed no discernible abnormalities in gross brain structures compared to those in WT brains. Measurement of cortical layer thickness in various regions, including the motor, somatosensory, visual, and auditory cortex, revealed no significant differences between FAM19A*5*-LacZ KI and WT mice. However, complete deletion of FAM19A*5* resulted in reduced brain weight and smaller brain structure^**4**^, suggesting its crucial role in early brain development. This notion is further supported by a recent study demonstrating that FAM19A*5* is predominantly expressed in germinal zones such as the ventricular zone and ganglionic eminence during the embryonic developmental stage^**3**^. These regions are pools of neural stem cells where division and differentiation into various types of cells occur. Therefore, FAM19A*5* likely plays a role in early brain development, and a partial knockout of FAM19A*5* likely has less of an impact on this process.

Furthermore, analysis of dendritic spine morphology in the motor and somatosensory cortices of FAM19A*5*-LacZ KI mice revealed an increase in the proportion of longer and thinner spines, indicative of a shift toward a more immature spine population. The spine density was not significantly altered in any of the analyzed brain regions. In contrast, a study by Huang et al., 2021 reported a significant decrease in spine density and a reduction in both mature and immature spine numbers in the CA1 region of the hippocampus in complete FAM19A*5* knockout mice^**4**^. This inconsistency in spine distribution may be attributable to a combination of early developmental issues, as discussed above, and technical limitations associated with analyzing low-resolution spine images and ambiguous classification criteria. For instance, the study reported the greatest number of mushroom spines, a type known for its widespread distribution in the brain^**18**^. Additionally, the quantification of even smaller spines, such as filopodia, was not included. These limitations may make it challenging to definitively assess the effects of FAM19A*5* knockout on the full spectrum of spine subtypes, potentially leading to underestimation of spine number changes.

FAM19A*5*-LacZ KI mice displayed hyperactivity, which is in accordance with the findings of a study by Huang et al.^**4**^. Hyperactivity is a characteristic feature of the neurodevelopmental disorder ADHD. There are few human cases in which patients with developmental defects are reported to have defects in chromosome 22, where FAM19A*5* is located^**7**,**8**^. Specifically, a microdeletion of chromosome 22, del22q13.32-q13.33, encompasses exon 4 of FAM19A*5* and is associated with clinical features such as hyperexcitability and aggression, similar to those observed in ADHD. This finding suggests a potential role for FAM19A*5* in neurodevelopment disorders. However, it is important to note that the microdeletion not only deletes the FAM19A*5* gene but also other neighboring genes, including SHANK3, which has been extensively studied in the context of ADHD^**19**,**20**^.

FAM19A*5*-LacZ KI mice showed increased activity, primarily in the peripheral region of the open field, with a greater tendency toward rearing behavior. Notably, the mice exhibited more supported rearing and consequently less unsupported rearing. Supported rearing is primarily associated with locomotor activity, whereas unsupported rearing is an exploratory behavior dependent on the hippocampus and can be indicative of a mouse’s emotional state^**14**^. This pattern suggested enhanced locomotor activity in FAM19A*5*-LacZ KI mice.

Furthermore, the FAM19A*5*-LacZ KI mice tended to exhibit less grooming behavior. Grooming is a measure of repetitive or compulsive-like behavior that occurs in individuals with neurodevelopmental disorders such as ASD^**21**^. Similarly, FAM19A*5*-LacZ KI mice displayed reduced marble burying behavior, a common test for repetitive actions^**22**^. This suggests a lack of repetitive behavior despite hyperactivity. The decrease in buried marble may be due to FAM19A*5*^LacZ/LacZ^ mice spending more time in other activities, such as supported rearing, leaving less time for digging.

Further behavioral analysis revealed the absence of anxiety/depressive-like behavior, normal spatial working memory, and normal recognition memory in FAM19A*5*-LacZ KI mice compared to WT littermates. However, a study on the FAM19A*5* knockout model by Huang et al. showed the presence of depressive-like behavior and impaired spatial memory^**4**^. The dynamic expression pattern of FAM19A*5*, with its potential role in early development^**3**^, suggested that complete knockout from conception might have a more profound impact than the partial reduction observed in our FAM19A*5*-LacZ KI model. This could explain the abnormal neurogenesis observed by Huang et al., which might in turn contribute to the learning and memory impairments reported in their study.

FAM19A*5* is highly expressed in rodent brain regions mediating fear learning and memory, including the amygdala and hippocampus^**23**^. FAM19A*5*-LacZ KI mice displayed impaired fear acquisition during the fear conditioning period. Consequently, analyses of both context-induced fear memory and tone-induced fear memory in mice were limited. The impaired fear acquisition in the mice might be due to their hyperactivity, but it might also be related to deficits in the innate fear response or sensory functions. Since all mice displayed similar vocalization and jumping behavior following unconditioned and conditioned stimuli during the fear acquisition period, their sensory function was likely intact. The unconditioned innate fear response test revealed a delayed response to TMT in FAM19A*5*-LacZ KI mice compared to WT mice. This finding suggested that the impaired fear acquisition in FAM19A*5*-LacZ KI mice may be related to a general deficit in their innate fear response.

Understanding the function of FAM19A*5* in the brain is crucial. Recent clinical studies revealed its genetic association with brain development-related symptoms such as ADHD and autism^8^, as well as neurodegenerative diseases, such as Alzheimer’s disease^24,25^. In this study, mice with partial FAM19A*5* reduction exhibited reduced body weight, abnormalities in dendritic spine maturation and structure, and hyperactive behavior. These findings strongly suggest that FAM19A*5* plays an important role in spine formation, synaptic transmission regulation, and normal brain development. Further supporting this notion, a recent study demonstrated that FAM19A*5* acts as a synaptolytic factor and that targeting it with antibodies in an Alzheimer’s disease could modify the disease course^5^.

In conclusion, this study employed FAM19A*5*-LacZ KI mice as a model for partial FAM19A*5* deficiency. Western blot analysis confirmed a reduction in FAM19A*5* protein levels in adult brains of these mice compared to wild type controls. Interestingly, FAM19A*5*-LacZ KI mice displayed lower body weight and reduced food intake, potentially linking FAM19A*5* to feeding behavior regulation in the hypothalamus. While gross brain structure appeared normal, further investigation into specific neuronal populations and their activity is warranted. Overall, these findings suggest a role for FAM19A*5* in body weight regulation and potentially early brain development, highlighting the need for further exploration of its function in the nervous system.

## MATERIALS AND METHODS

### Animals

All mice were housed under temperature-controlled (22–23°C) conditions with a 12-h light/12-h dark cycle (lights on at 8:00 am). The mice were given ad libitum access to standard chow and water. All animal experiments were designed to use the fewest mice possible, and anesthesia was administered before sacrificing the mice. All animal procedures were approved by the Institutional Animal Care and Use Committee of Korea University (KOREA-2016-0091-C3).

Homozygous FAM19A*5*-LacZ KI mice were obtained by mating heterozygous male and female FAM19A*5*-LacZ KI mice. The WT littermates were used as the control group. Body weights were measured weekly starting from four weeks to ten weeks after birth. Brain sizes were measured during brain sampling using a digital Vernier caliper. The daily food intake of six-week-old male mice was measured for 15 days. For the behavior test, ten twelve-week-old male mice were used unless otherwise mentioned.

### Measurement of cortical thickness

Nissl-stained sections were imaged using a slide scanner (AxioScan Z1, Zeiss). Different brain regions were defined using the mouse brain atlas (motor cortex, bregma 1.34∼0.98 mm; somatosensory cortex, bregma -1.22∼-1.*5*8 mm; visual cortex, bregma -2.*5*4∼2.92 mm; and auditory cortex, bregma - 2.*5*4∼2.92 mm). For cortical layer measurements, different layers were distinguished based on the cellular morphology and arrangement. Three nonadjacent Nissl-stained brain sections at intervals of 120 μm from each genotype (n=*5*) were analyzed. The average cortical thickness and layer thickness were calculated from three independent linear measurements using ZEN software (Zeiss).

### Golgi staining and dendritic spine analysis

Ten-week-old mouse brains were subjected to spine analysis. Golgi staining was performed using the FD Rapid GolgiStain Kit (FD NeuroTechnologies) following the manufacturer’s protocol. Briefly, mouse brains were isolated and immersed in impregnation solution (a mixture of solutions A and B) for two weeks in the dark at room temperature. Next, the brains were transferred to Solution C and stored at room temperature in the dark. Coronal sections 100 μm thick were obtained using a vibratome and transferred to gelatin-coated slides (Lab Scientific). The sectioning procedure was carried out in Solution C to prevent cracking of the sections. Solution C was blotted completely from the slide, and the sections were then air-dried at room temperature overnight, washed with DW three times every *5* min and subjected to working solution (a ratio of 1:1:2 of solution D:E:DW). After washing in DW 3 times every *5* min, the sections were dehydrated in increasing concentrations of ethanol (50%, 75%, 90% and 100%) for *5* min each, cleared in xylene and coverslipped using Permount solution. Z-stack images (0.5 μm intervals) of the basal and apical dendrites of approximately 5-7 pyramidal neurons in layer 2/3 and layer *5* of the motor cortex and somatosensory cortex were acquired using a 60X objective with a confocal microscope (TCS SP8, Leica). Spines from secondary basal and apical dendrite segments of at least 20 μm in length were selected for quantitative analysis. Approximately *5* dendritic segments from each neuron were analyzed. The number of spines and the width and length of an individual spine were manually measured as previously described^***10***^. The spine density was expressed as the number of spines/10 μm of dendrite. The length-to-width ratio (LWR) was calculated as a measure of spine morphology.

### Western blot analysis

Mice were decapitated, and brains were isolated immediately and chilled in ice-cold PBS. The brain tissues were homogenized in lysis buffer (10 mM Tris-HCl, pH 7.5, 500 mM NaCl, *5* mM MgCl2) containing protease inhibitor cocktail. The homogenate was centrifuged for 10 min at 15,000 rpm, after which the supernatant was obtained. The protein concentration of the supernatant was determined using a Bradford protein assay kit (Bio-Rad). The brain lysates were then denatured in SDS sample buffer and separated by SDS-PAGE. The proteins were transferred to a polyvinylidene difluoride (PVDF) membrane and blocked with a 5% skim milk solution for 30 min at room temperature. The membrane was incubated with a FAM19A*5* 3-2-HRP-conjugated antibody (20 µg) overnight at 4°C. After washing with TBST 3 times every 10 min, the signals were detected using an enhanced chemiluminescence (ECL) assay kit (Thermo Scientific). β-Actin was used to normalize the levels of protein detected.

### Open field test

The open field test (OFT) was used to assess exploratory behavior and general locomotor activity. Mice were placed in a white opaque open field box (40 × 40 × 40 cm) and allowed to explore freely for 10 min. The field was divided into two zones, the central and peripheral zones. The zone 10 cm from the edge of the field was defined as the center zone. The total distance traveled, mean speed of movement and total time spent in the center of the field were analyzed using ANY-Maze software (Stoelting). In addition, ethological parameters such as rearing and grooming were analyzed manually.

### Elevated plus maze test

The elevated plus maze test (EPMT) was used to measure anxiety-like behavior. The maze consisted of two oppositely positioned open arms (5 X 30 cm) and an oppositely positioned closed arm (5 X 30 cm) enclosed by a 20 cm high wall placed 50 cm above the ground. Mice were placed in the center of the maze facing the open arm and allowed to freely explore the maze for 10 min. The total time spent in the open arms, total distance traveled and mean speed of movement in the maze were analyzed using ANY-Maze software (Stoelting). Arm entry was considered only when all four paws were inside the defined zone.

### Novel object recognition test

The novel object recognition (NOR) test was used to assess recognition memory. The test was conducted in an open field box (40 X 40 X 40 cm) with two distinct objects (Lego block and T-75 flasks filled with sand). Mice were habituated to the open field without any objects for 10 min one day before the training session. On the next day, during the training phase, the mice were allowed to freely explore the field containing two identical objects for 10 min. The test phase was performed after 6 h for short-term memory and after 24 h for long-term memory. In the test session, one of the objects was replaced with a novel object, and the mice were placed in the field for 10 min. The time spent exploring each object was analyzed using ANY-Maze software (Stoelting). Preference for the novel object was defined as the time spent exploring the novel object divided by the total time spent exploring both objects.

### Y-maze test

***The*** Y-maze test was used to measure spatial learning memory. Mice were placed in the center of the Y-shaped maze with three identical arms (30 X *5* X 20 cm), and movement was recorded for *5* min. The total number of arm entries and sequence of entries were analyzed using ANY-Maze software (Stoelting). Entry occurs when all four limbs are within the arm. Spontaneous alteration was defined as the number of triads (ABC, ACB, BAC, BCA, CAB, CBA) divided by the total number of arm entries minus 2 multiplied by 100.

### Marble burying test

The marble burying test was used to study repetitive behavior and anxiety-like behavior. A cage containing a total of 20 marbles arranged on the surface of *5* cm thick bedding in *5* rows of 4 marbles was employed. Mice were placed in cages containing marbles and left undisturbed for 30 minutes. The total number of marbles buried was counted and plotted as a graph. Marbles were considered buried if 2/3rd or more of the surface area was covered by bedding.

### Nestlet shredding test

The nestlet shredding test was used as a measure of repetitive compulsive-like behavior. A nestlet was weighed and placed in the cage with 0.5 cm thick bedding. Mice were placed into the cage for 30 min without any disturbance. Food and water were also provided during the test period. After completion of the test, the nestlets were allowed to dry overnight to avoid moisture. The next day, the unshreded nestlets were weighed, and the percentage of shredded nestlets was plotted.

### Rotarod test

The rotarod test was performed to assess motor learning and coordination. The mice were placed on a rotating rod at increasing speeds (from 4 to 40 rpm) for *5* min. The test was carried out for *5* consecutive days, and the latency to fall and the time it took the mice to fall off the rotating rod were recorded. The mice were subjected to three trials per day, with a maximum of 300 s for each trial and an interval of at least 15 min between trials.

### Hanging wire test

The hanging wire test was used to measure muscle strength, mainly on the forelimbs and hindlimbs. Mice were placed on top of wire mesh, which was then inverted, and the mice were allowed to hang for a maximum of 300 s. Five repetitive trials were performed in which the first 2 trials involved acclimatization, and the latency to fall during the remaining 3 trials was recorded, the highest of which were plotted.

### Tail suspension test

The tail suspension test (TST) was used to measure depressive-like behavior. Mice were suspended above the ground by the tail with the aid of adhesive tape for 6 min. The movement of the mice was monitored, and immobility was measured using ANY-Maze software (Stoelting). Immobility was defined as the time at which the mice stayed without body movement.

### Pavlovian fear conditioning test

The Pavlovian fear conditioning test was conducted to study fear response memory, as previously described^***26***^. Briefly, the mice were habituated to the fear conditioning chamber for 10 min without any stimulus. The next day, a training session was performed in which 7 repetitive trials, each consisting of a tone (*5* kHz, 70 dB, 30 s) that terminated with a foot shock (0.7 mA, 2 s), were delivered. The mice were returned to their home cages, and the test session was conducted the following day. For the contextual memory test, the mice were placed in the same chamber for *5* min without tone or electric shock. The freezing time was analyzed using ANY-Maze software (Stoelting). For the auditory memory test, the mice were placed in a distinct context and exposed to 3 tones without foot shocks with an intertrial interval of 90 s after *5* min of acclimatization. The freezing time during each tone was analyzed using ANY-Maze software (Stoelting), and the average freezing time was calculated.

### Unconditioned innate fear response test

An unconditional innate fear response test was performed to evaluate innate fear, as previously described (Yun et al., 2019). Briefly, mice were placed in a chamber in which 30 µl of 2,*5*-dihydro-2,4,*5*-trimethylthiazoline (TMT), a synthetic component found in fox feces, was added. The video was recorded for 1*5* min, and TMT-induced freezing was analyzed using ANY-Maze software (Stoelting). The percentage of freezing time for every 3 min was plotted.

### Statistical analysis

All the statistical analyses were performed using GraphPad Prism *5* software (GraphPad Software, Inc., La Jolla, CA). The data are shown as the means ± SEMs. Statistical significance was determined using two-tailed unpaired Student’s t test. For multiple comparisons, two-way ANOVA followed by the Bonferroni post hoc correction was performed. The criterion for statistical significance was set at a p value less than 0.0*5*.

## DATA AND MATERIALS AVAILABILITY

The raw data and genetic constructs are available upon request to the authors.

