## Supplementary Materials for "Partial FAM19A5 Deficiency in Mice Leads to Disrupted Spine Maturation, Hyperactivity, and an Altered Fear Response"

### Supplementary Information

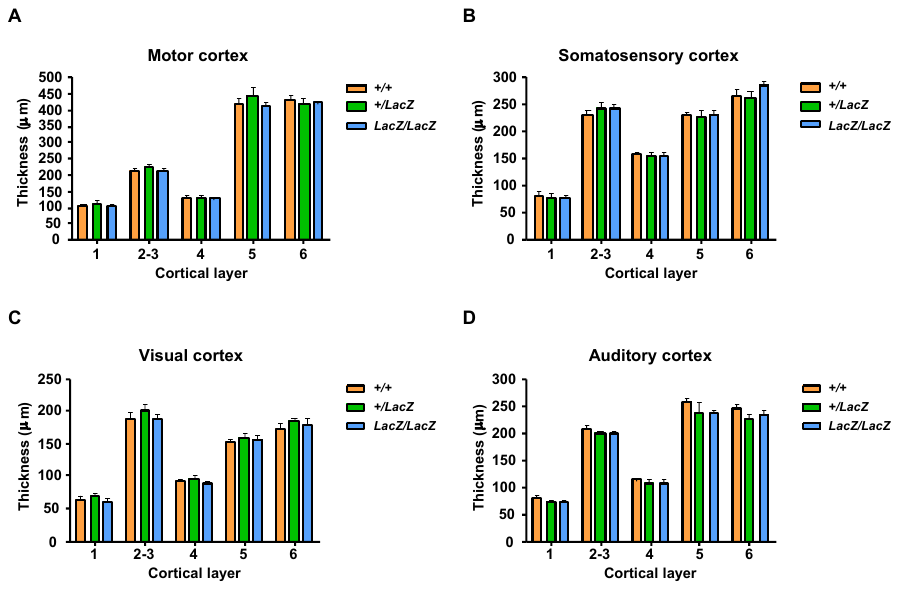

**Supplementary Fig. 1 |** **Cortical layer thickness of adult FAM19A5-LacZ KI mouse** The cerebral cortical layer thickness in (A) motor, (B) somatosensory, (C) visual, and (D) auditory cortex in FAM19A5-LacZ KI and FAM19A5^+/+^ mice brains (n=5). Data are presented as the mean ± SEM.

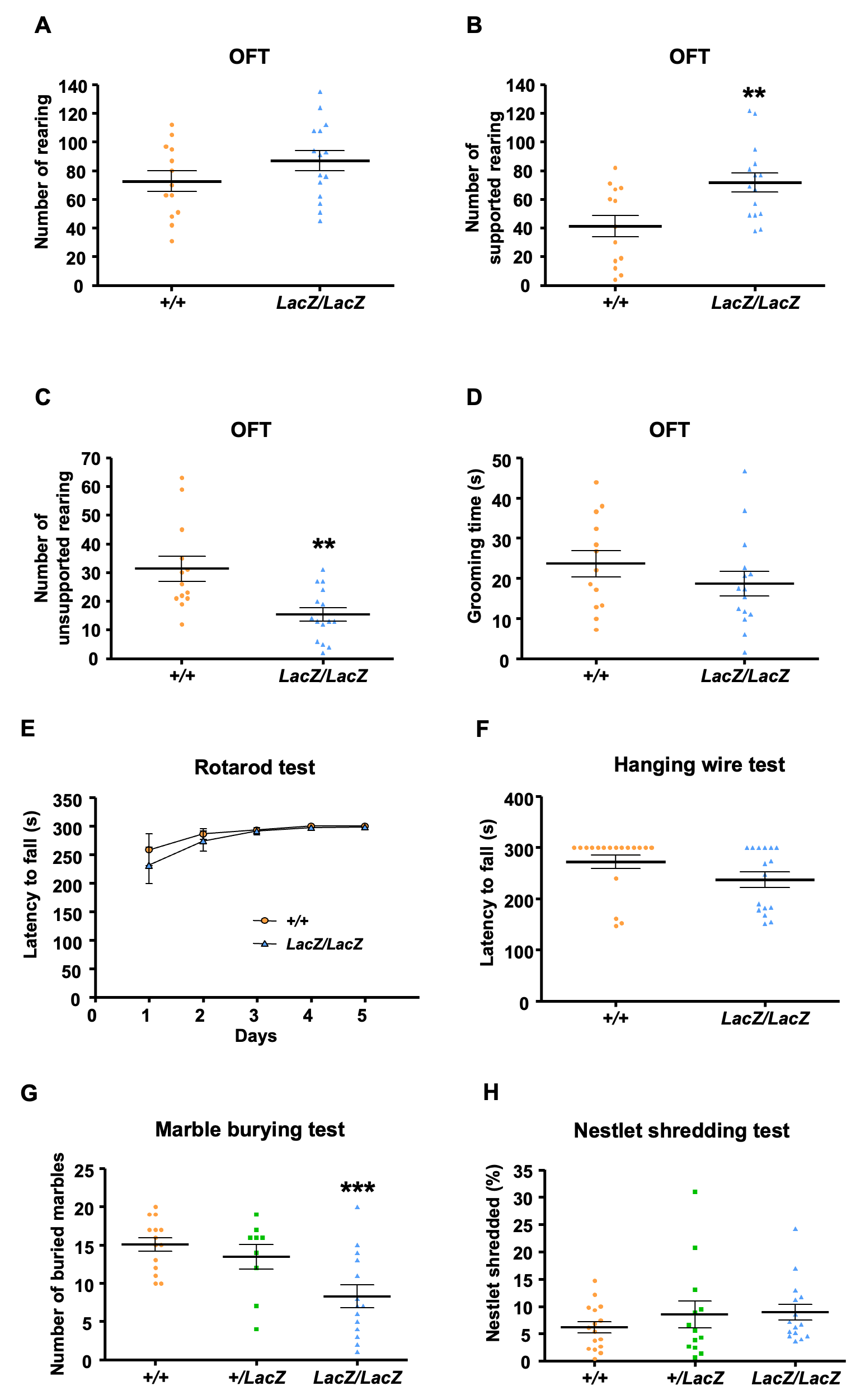

**Supplementary Fig. 2 |** **Rearing and grooming behavior in FAM19A5-LacZ KI mice** (A) Total number of rearing, (B) supported rearing, (C) unsupported rearing, and (D) total grooming time during 10 min of exploration period in OFT. FAM19A5^+/+,^ n=13 and FAM19A5^LacZ/LacZ^, n=15. Data are presented as the mean ± SEM. **P<0.01 vs. FAM19A5+/+. **(**E) Latency to fall in a rotating rod during 3 trials per day for 5 consecutive days (FAM19A5^+/+^, n=9 and FAM19A5^LacZ/LacZ^, n=7). (F) Latency to fall in a hanging wire test in 300 s long 3 consecutive trials with at least 15 min intertrial interval (FAM19A5^+/+^, n=18 and FAM19A5^LacZ/LacZ^, n=16). (G) Number of marble buried during 30 min of exploration time (FAM19A5^+/+^, n=14; FAM19A5^+/LacZ^, n=9; FAM19A5^LacZ/LacZ^, n=14). (H) Percentage of shredded nestlet during 30 min of observation time (FAM19A5^+/+^, n=16; FAM19A5^+/LacZ^, n=13; FAM19A5^LacZ/LacZ^, n=15). Data are presented as the mean ± SEM. ***P<0.001 vs. FAM19A5^+/+^.

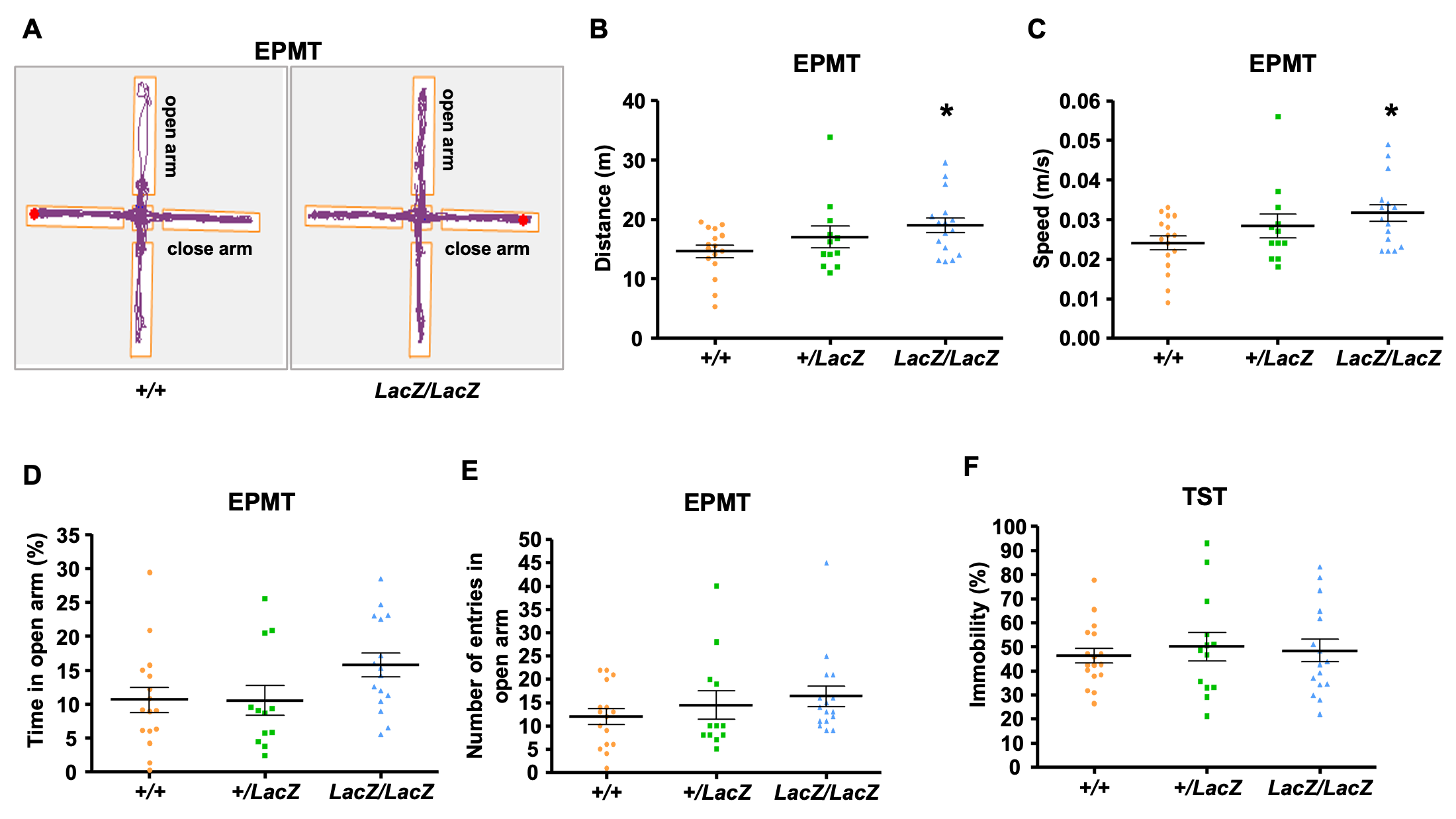

**Supplementary Fig. 3 |** **The absence of anxiety and depressive-like behavior in FAM19A5-LacZ KI mice** (A) Representative track plot of movements in elevated plus maze during 15 min of exploration time in FAM19A5^LacZ/LacZ^ and FAM19A5^+/+^ littermates. (B and C) Total distance traveled and mean speed of movement in elevated plus maze, respectively. (D and E) Percentage of time spent and total number of entries in open arm of the maze, respectively. (F) Percentage of immobile time during 5 min long TST. FAM19A5^+/+^, n=16; FAM19A5^+/LacZ^, n=12 and FAM19A5^LacZ/LacZ^, n=16. Data are presented as the mean ± SEM. *P<0.05 and **P<0.01 vs. FAM19A5^+/+^.

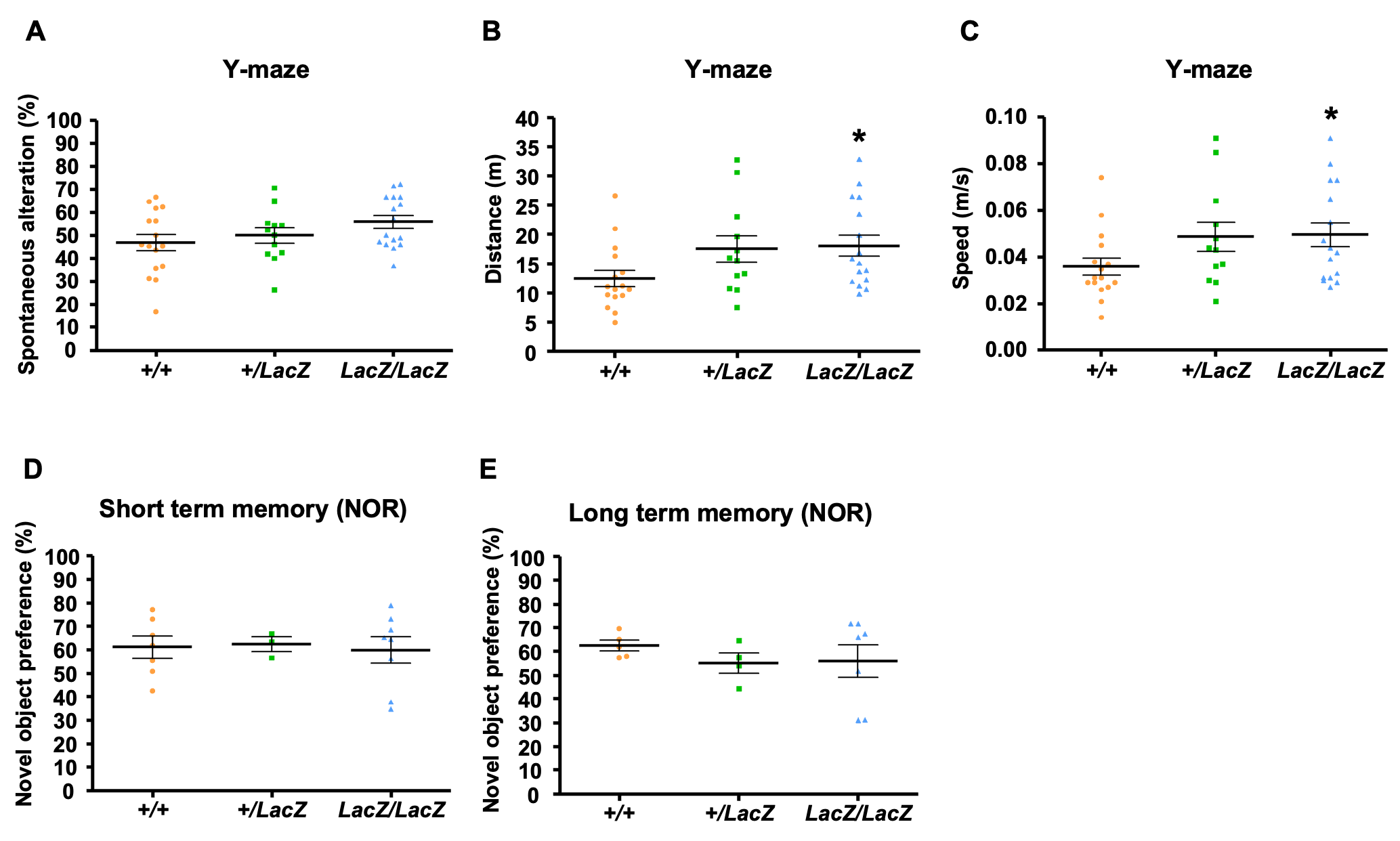

**Supplementary Fig. 4 |** **Normal learning and memory function in FAM19A5-LacZ KI mice** (A-C) Total distance traveled, speed of motion and percentage of spontaneous alteration in Y-maze arms during 5 min of exploration period in FAM19A5^+/LacZ^, n=12; FAM19A5^LacZ/LacZ^, n=16 and FAM19A5^+/+,^ n=16. (D) Percentage of preference to novel object during 10 min exploratory time in NOR test after 6 h for short term memory and (E) after 24 h for long term memory. FAM19A5^+/+^, n=7; FAM19A5^+/LacZ^, n=3; FAM19A5^LacZ/LacZ^, n=8 for short term memory and FAM19A5^+/+^, n=5; FAM19A5^+/LacZ^, n=4; FAM19A5^LacZ/LacZ^, n=7 for long term memory. Data are presented as the mean ± SEM. *P<0.05 vs. FAM19A5^+/+^.
